# *plinkQC*: An Integrated Tool for Ancestry Inference, Sample Selection, and Quality Control in Population Genetics

**DOI:** 10.1101/2025.11.25.690541

**Authors:** Maha Syed, Caroline Walter, Hannah V. Meyer

## Abstract

**Motivation:** Population genetic analyses rely on high quality datasets that pass rigorous controls for sample and marker quality. Many analyses also require additional processing including identification of ancestry and sample relatedness. A software package that addresses all these common, yet crucial tasks is missing.

**Results:** We have developed *plinkQC*, an R/CRAN package that combines these functionalities into a single software package with detailed vignettes for example applications. *plinkQC* determines the ancestry of study samples via a pre-trained random forest classifier that reaches 98% performance accuracy with just 5% of marker overlap between reference and user data. To obtain the maximal set of unrelated study samples, we developed a graph-based pruning method, taking both relationship estimates and sample quality into account. We demonstrate optimal sample selection on the 1000 Genomes project, where we retain an additional 71 samples compared to publicly available exclusion lists. Finally, *plinkQC* bundles these results together with per-individual and per-marker quality control checks into three simple functions and returns both the quality controlled data set and quality control report about each step of the analysis.

**Availability:** *plinkQC* is available as an R/CRAN package. The documentation and code are available on github: https://meyer-lab-cshl.github.io/plinkQC/ and https://github.com/meyer-lab-cshl/plinkQC_manuscript.

## Introduction

Genetic association tests such as genome-wide association studies (GWAS) and quantitative trait loci (QTL) mapping are vital to our understanding of genetic risk factors and underlying biological mechanisms of physiological traits and diseases^1–3^. For robust and reproducible association results, rigorous quality control (QC) of the genotype data by removing low-quality samples and genetic markers is crucial. QC is commonly based on summary statistics of the genotypes, and is conducted on a genetic marker (most commonly single nucleotide polymorphism; SNP) and sample level^4^. Genetic Marker QC includes removing markers with high missingness, low frequency and those not in Hardy-Weinberg equilibrium due to presumed genotyping errors. Sample-level quality controls include tests for sample swaps, often by checking if the predicted sex matches the assigned sex, the rate of missing markers, and the observed within-sample heterozygosity^4^. In addition, analyses often require controlling for a shared genetic background and/or direct relatedness within the study cohort. Depending on the purpose and assumptions of the downstream analysis, some or all of these filters may be applied to the data.

Currently, rigorous genotype QC relies on the use of multiple tools to complete different tasks. For example, *King*^5^ is commonly used to remove related samples, while Admixture^6^ is used to determine the ancestry of samples. In addition, there is no tool that provides a standardized QC framework, with an accompanying report and direct output of the processed dataset with all filters applied. Here, we describe *plinkQC*, an efficient and all-in-one R/CRAN package to perform QC, ancestry identification, and selection of the maximally unrelated sample set (Figure 1). It takes in variant based genotyping data and returns a post-QC dataset of high-quality markers and samples, with predicted genomic ancestry and relatedness. For the latter, *plinkQC* implements i) a classifier that predicts the genomic ancestry of human samples and ii) graph-based relatedness pruning taking sample quality metrics into account. For QC, *plinkQC* computes underlying statistics with the commonly used command-line program *plink* and visualizes results in a QC report. In summary, *plinkQC* bundles important functions to make genomic QC more efficient and accessible.

**Figure 1.**
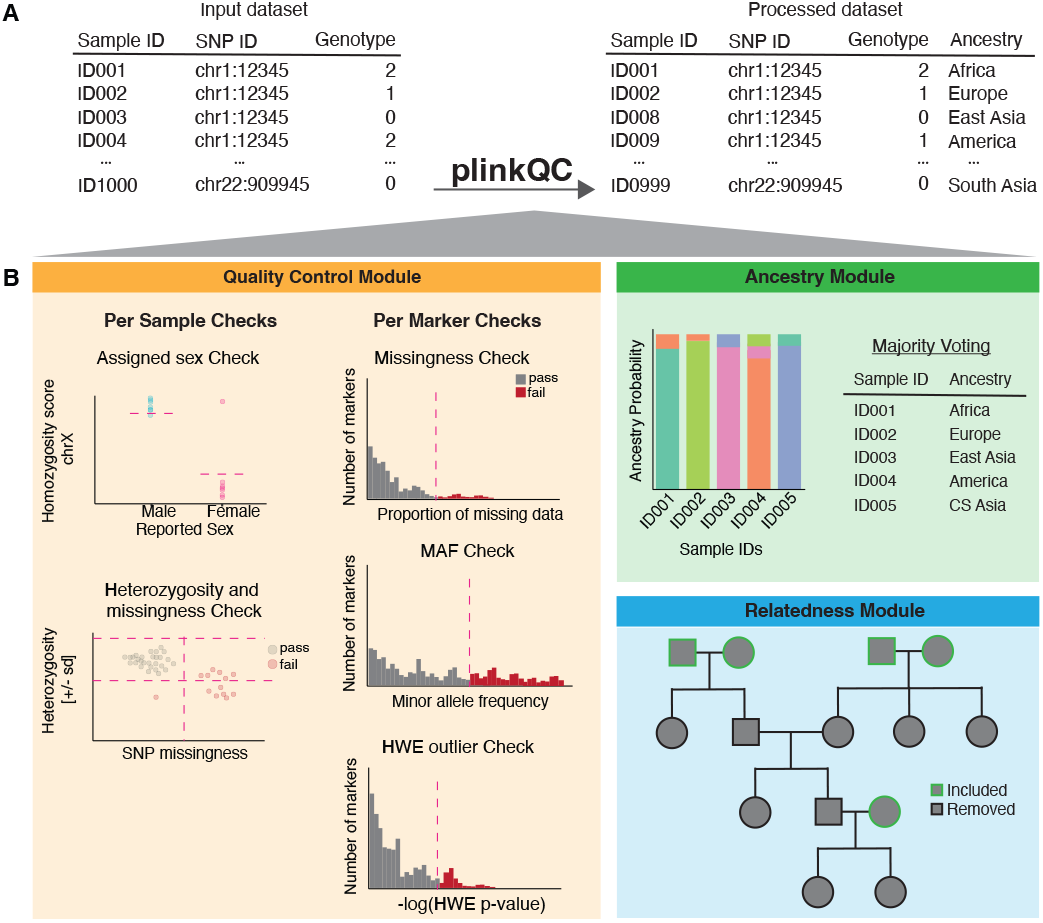
Standardized and modular genotype processing with *plinkQC*. A) *plinkQC* takes in marker-based genotype data as input and returns the processed dataset based on user-set modules and filters. B) Quality control module: per-sample and per-marker checks for missing data, sample swaps and marker properties. Ancestry module: probabilistic and classification-based ancestry prediction. Relatedness module: relatedness identification and construction of maximal unrelated and high QC sample set.

### Workflow

*PlinkQC* takes in variant-based genotype calls in the common *plink* format with files containing sample information (.*fam* files), genotype information including genomic position (.*bim* files) and the actual compressed genotypes (binary.*bed* files) (visualized as human-readable tables in Figure 1A). QC is conducted on a per-marker and per-sample level and users can choose from a set of QC filters (Figure 1B). *plinkQC* first computes and visualizes the QC statistics, providing the user with the choice of thresholding for subsequent application of marker and sample removal that fail QC.

#### Per-marker QC

The following three per-marker QC steps are recommended: i) removal of genetic markers with high missingness rate as they might not have been genotyped properly; ii) removal of genetic markers with a low minor allele frequency (or minor allele count; as specified by the user); they are often removed as downstream analyses would be underpowered and iii) removal of genetic markers with low p-value (high −log_10_(p-value) in the test for Hardy-Weinberg equilibrium (HWE). This step is included as the strong deviation from HWE is assumed to reflect genotype calling errors. However, there could also be true signal captured in these markers and to avoid excluding biologically relevant markers under selection in case-control studies, often only the control samples are used to determine markers to remove.

#### Per-sample markers

The following three per-sample QC steps are recommended in each analysis: i) prediction of the biological sex of each sample and comparison to the recorded sex to identify potential sample mislabeling or data entry errors; ii) removal of samples with high missingness rate to identify poor quality samples and iii) removal of samples with outlying heterozygosity rate to flag samples that are potentially inbred or are contaminated with external DNA. In addition, downstream analyses might require assessing correlations between samples. Genomic data can be correlated between samples because of direct relatedness and/or broader population substructure. To support this, *plinkQC* includes tools for identifying related individuals and inferring genomic ancestry — key steps for controlling confounding effects and enabling robust downstream analysis. Users can choose which if any of these checks they would like to run.

### Reference Dataset

To develop the ancestry and relatedness modules, we relied on well-annotated human datasets, capturing a broad range of genetic backgrounds. The harmonized dataset of samples from the 1000 Genomes^7^ and Human Genome Diversity Project (HGDP)^8^ served as an ideal reference data set^9^, with 4,151 individuals spanning seven continental ancestry groups.

In addition to representing a wide range of geographic and ancestral populations, the 1000 Genomes dataset also includes known familial relationships, providing a ground truth dataset for the development of our relatedness estimation module. After subsetting for samples from the 1000 Genomes project, the dataset contained 3,202 samples. These samples were split into 2,504 unrelated individuals and 698 individuals related to those in the prior dataset. From the 1000 Genomes pedigree file, there are 1,799 samples labeled as having first degree relatives, and an additional 27 samples labeled as having second degree relatives.

For the development of the ancestry prediction module, we used all unrelated individuals of the combined 1000 Genomes and HGDP passing quality control. We combined the South Asia label from the 1000 Genomes dataset with the Central South Asia label from the HGDP dataset due to an overlap in the geographical regions. The final dataset contained 3,379 samples across six continental ancestry groups: Africa, Central South Asia (CS Asia), Europe, admixed America, East Asian and Middle Eastern (Figure 2A).

**Figure 2.**
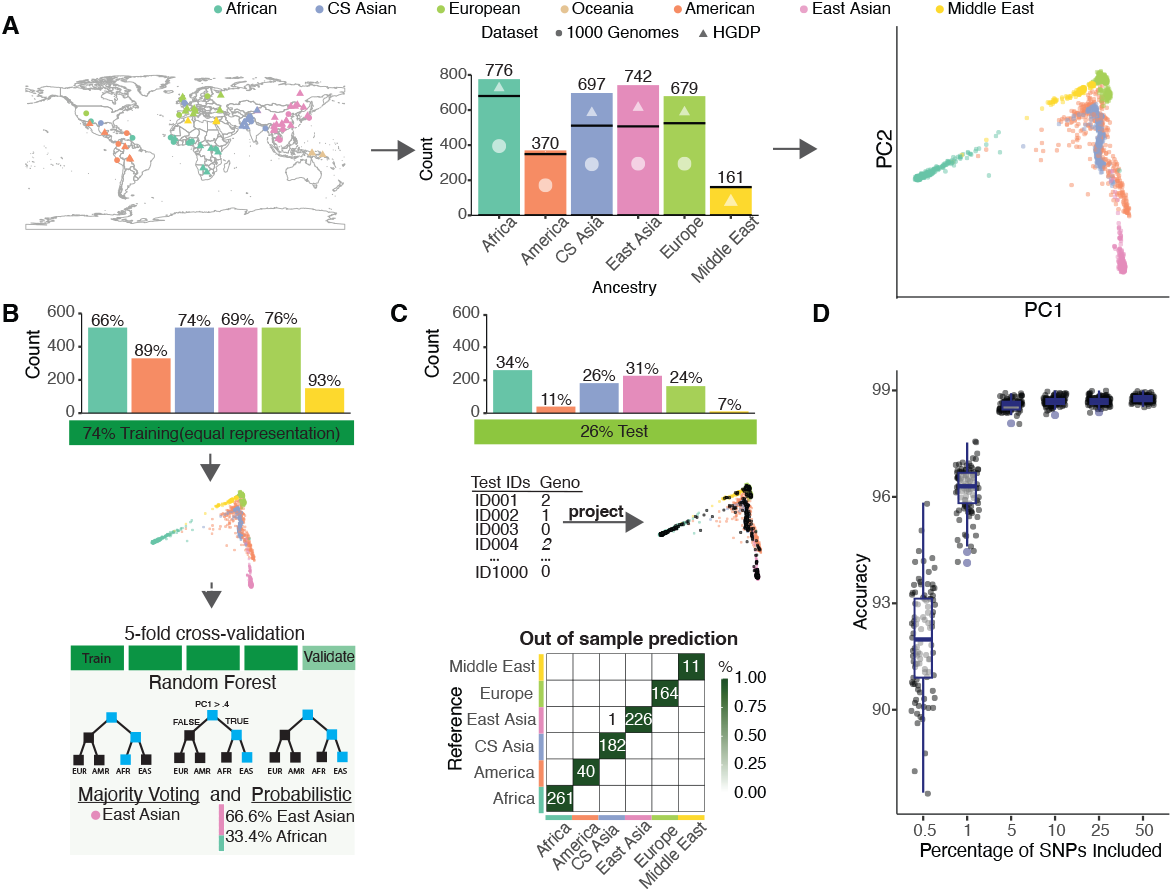
Ancestry Identification with *plinkQC*. A) Geographical distribution and sample counts of the full harmonized reference dataset consisting of 4,151 samples from combined the 1000 Genomes (3,202 samples) and Human Genome Diversity Project datasets (948 samples). Principal component (PC) computed on the post-QC dataset of 3,436 individuals with 229,020 independent SNPs (LD *r*^2^ < 0.2. B) Schematic of training data and random forest classifier taking PCs as features for ancestry prediction as either a probability vector or classification. C) Ancestry classification of held-out test data by projecting test samples’ SNPs into PC embeddings based on training set. Classification accuracy is 99.9%. Zeros are not marked and the colors are row normalized. D) Accuracy of the algorithm on the held out test data with random downsampling of SNPs (n=100 trials).

### Ancestry Identification

Population stratification can lead to spurious results in genetic association studies where associated variants are related to a group membership instead of the phenotype^10^. It occurs in data where subgroups display increased genetic correlation, which, in human data, is observed in individuals of common genomic ancestral backgrounds.

Without a sample pool that reflects the makeup of the population, the insights from genetic association studies are limited. Risk variants and effect sizes found in one ancestral group do not always correlate between ancestries^11^, which may be due to differences in linkage disequilibrium (LD) patterns^12^ or varying pleiotropic effects of different genetic backgrounds. Additionally, ancestral groups may have different frequencies of risk alleles, and successfully identifying and including them in association studies enables novel risk allele identification.^13,14^. Thus, designing association tests that use all sample ancestries is critical to fully understand the effects of human genetic variation.

To account for population stratification in genetic association studies, many methods have implemented principal components (PCs) of the sample genotypes. For cohorts composed of multiple ancestries, PCs can be used to define and divide cohorts into ancestry-specific subgroups for independent analyses and potential meta-analyses^15,16^. Even within a cohort composed of one major continental ancestry, PCs may be used to exclude outliers^17^ or capture small scale differences that can be accounted for as covariates in the model of choice^18^.

Historically, the majority of human GWAS have been conducted on individuals from European descent^19^. To reduce genetic heterogeneity, many of these studies removed non-European ancestral samples from their analysis^17,20^. However, with growing efforts to expand genomic data acquisition and genetic association studies beyond European ancestries, we need to be able to identify and retain multi-ancestral samples in the analysis.^21–25^.

Here, we trained a random forest classifier on the combined 1000 Genomes HGDP reference panel. It takes projected sample PCs as features and returns estimated ancestry as either a probability vector or a classification (Figure 2B, barplot). Our classifier achieves a 99.9% accuracy rate on the held-out test data, with one sample with Africa ancestry misclassified as having America ancestry (Figure 2D). To evaluate the robustness of PC-based random forest classifier under conditions of incomplete genetic data, we simulated data with reduced marker overlap between user-provided input and the reference dataset. With just 5% of the SNPs shared between the reference and input data, the mean accuracy was 98.6%, close to the 99.9% accuracy with using the full set of markers (Figure 2F).

As an external evaluation to assess how well our ancestry inference generalizes to a new dataset, we used SNP genotypes of 100 Fulani individuals from the Gambian Genome Variation Project^23^ (Figure 3A). In addition to external validation, we also used this analysis to benchmark our ancestry inference method with two existing tools: ADMIXTURE^6^ and SNVStory^26^. ADMIXTURE is a commonly used program for estimating an individual’s ancestry proportions. Setting up ADMIXTURE is a two-step process. First, it is run unsupervised on the reference dataset and individuals with high levels of admixture statistic are removed for subsequent training in supervised mode on the smaller dataset. The classifier built in supervised mode is then used for predicting sample ancestry. As the reference dataset SNPs should match the study dataset, it is often necessary to retrain the model. *SNVstory* uses a support vector machine directly on SNPs for ancestry identification. To set up *SNVstory*, we installed Docker Desktop and executed the software within the provided Docker Container. *plinkQC*, ADMIXTURE and *SNVstory* all predict 100% African ancestry. *SNVstory* has a much longer runtime than *plinkQC* and ADMIXTURE (Figure 3B). In addition to the upfront set-up time, ADMIXTURE’s total time for the two-step training and projecting the 100 Fulani samples takes longer than *plinkQC*’s ancestry identification (time breakdown for each step given in Supplementary Fig 2). Overall, *plinkQC*’s ancestry identification module is able to correctly and efficiently identify ancestries of external data without additional processing or classifier retraining.

**Figure 3.**
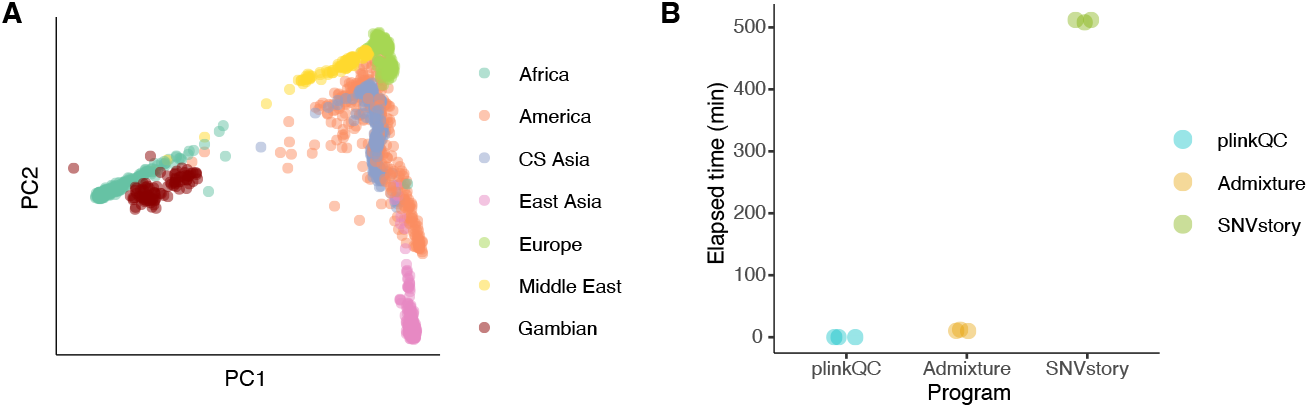
Validation and Benchmarking on Gambian Genome Variation Project. (A) Genotypes (14,095 SNPs) of 100 Fulani individuals from the Gambian Genome Variation Project projected into PC1 and PC2 of combined 1000 Genomes and HGDP reference panel. (B) Empirical runtime analyses for estimating ancestries of 100 Fulani individuals with *plinkQC*, ADMIXTURE, and *SNVstory*. The elapsed time to run ADMIXTURE is the combined time of unsupervised and supervised training and classification.

### Relatedness

Genetic association analyses and heritability estimation may require removal of related individuals. However, the decision of which individuals to remove can influence the overall sample size. For instance, in a dataset containing two parents and one child, it may be more advantageous to retain both parents and exclude the child instead of retaining just the child. Within *plinkQC*, we developed a method to infer the largest independent vertex set of samples within a sample-relatedness graph to retain the maximum number of samples.

The approach begins by estimating identity-by-descent (IBD) between all pairs of samples. Predicted IBD is calculated by the proportions of the genome that is shared between two individuals^27,28^. If the pair of samples have a predicted IBD above a user-defined threshold, the samples are considered related. From the individuals labeled as related, we divide the sample pool into subgroups of related families, from which we construct family-specific vertex sets. Based on these, we calculate the maximum independent vertex set between individuals. If there are multiple options, for example one parent and two offspring, *plinkQC* chooses samples based on higher QC metrics.

To test the functionality of the relatedness pruning, we used the 1000 Genomes dataset, which contains 2,504 unrelated individuals from the primary release and an additional 698 individuals related to the first sample set. The release of these two datasets was separated by time and the 2,504 individuals labeled as unrelated do not necessarily build the maximal unrelated dataset out of the total of 3,202 individuals. Here, we both address the question of the maximal unrelated set in the 1000 Genomes dataset and benchmark our approach to *King*^5^, a tool for detecting and filtering related individuals. In addition, since *plinkQC* and *King* use different metrics to estimate relationship, we also compared the *plinkQC* relatedness filter using *King* generated kinship scores as input (Figure 4C).

**Figure 4.**
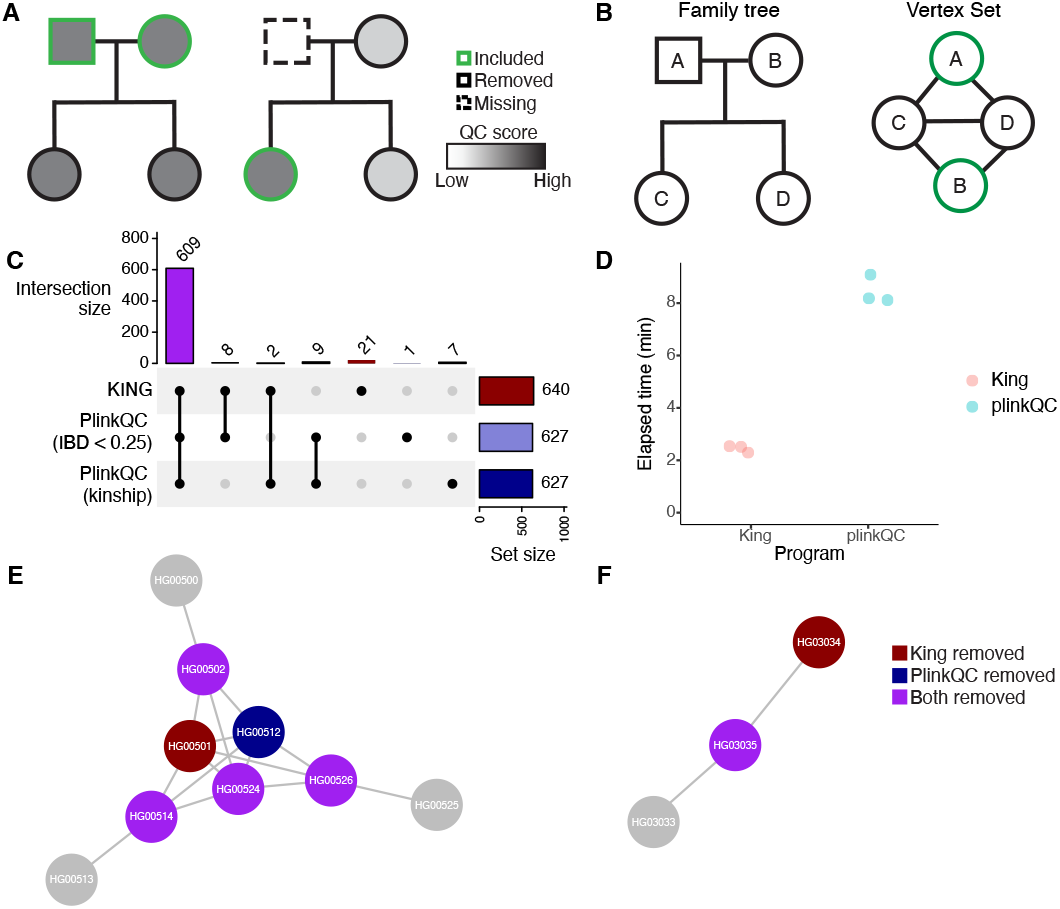
Relatedness identification and selection of unrelated individuals. A) Schematic showing sample selection with pruning out related individuals based on retaining the maximum number of individuals with high QC scores. B) Schematic of a four person family tree and its corresponing vertex set C) Samples from the 1000 Genomes dataset that are pruned out by *King, plinkQC* using the built in IBD scores, and *plinkQC* using *King* kinship scores as a relatedness matrix. The bars on the top show the intersection of samples that are removed and the bars on the right represent the total samples removed within each group. D) User elapsed time taken to run *King* and *plinkQC* on the 1000 Genomes Dataset. Points are jittered for readability. E) and F) An independent vertex set of a family structure and trio found within the 1000 Genomes. A line between samples indicates a kinship score of above the *King* threshold for second degree relatives of 0.0884. Colors represent samples removed by *King, plinkQC* (kinship), or both.

The majority of samples removed by *King* and *plinkQC* (with either relatedness estimate) are shared (609 samples; Figure 4, intersection size). Looking closely at samples that are removed uniquely by any method, we see that they fall into two categories. They are either a pair of individuals, where removal of either individual reduces the relatedness in the sample set by same amount (e.g. removal of two out of three siblings; Figure 4E) or *King* removes superfluous individuals e.g. two individuals from a trio when only one needs be removed (Figure 4F). Overall, *plinkQC* with either IBD or *King*-based kinship as a relatedness estimate removes a consistently lower number of individuals than *King*. We used the 1000 Genomes pedigree file to determine how well these metrics match the annotated relatedness. We find that both algorithms pruned all family structures indicated as related. In addition, *King*- and IBD-based relatedness estimates uncovered related samples i.e. *King* kinship and IBD scores above the respective thresholds, that had no labeled relatives in the pedigree files that were subsequently removed in the relatedness filter. In addition, we see there are 4 individuals labeled as secondary degree related which were not removed by *plinkQC* and 3 that were not removed by *King*. However, based on both IBD and kinship scores these individual do not pass a threshold of secondary degree relationship (Supplementary Table 1). In summary, we identified that the maximal set of individuals without first and second degree relatedness in the 1000 Genomes dataset is 2,575, increasing the sample size by 71 individuals.

## Conclusion

We have introduced *plinkQC*, an open source R/CRAN package for genotype QC, relatedness, and ancestry estimation. Its QC module contains functions to conduct user-defined per-marker and per-sample quality control checks and return both the cleaned dataset as well as an automated QC report. Importantly, *plinkQC* offers additional functionality, with its relatedness and ancestry modules identifying maximally unrelated sample sets and ancestries of the study population. Combining these functions into single, software package with detailed vignettes for example applications, we provide an easy to install and to use solution for crucial, yet common tasks prior to genetic data analyses.

Custom random forest implementations have successfully been used for ancestry classification based on SNP genotypes^9,25^. Here, we trained a random forest classifier on a large human reference panel, which provides ancestry estimation in minutes without study-specific retraining of the classifier. Further, we showed that classification accuracy remains excellent even if the overlap between SNPs in the study cohort and the training data is as little as 5%. *plinkQC*’s classification performance on external validation data is consistent with prior methods *SNVstory* and ADMIXTURE while improving both ease of setup and runtime. For ADMIXTURE, commonly multiple training runs using different initiation seeds have to be run to find a converging classifier. Additionally, users must include their own reference dataset, where reference and study dataset must contain the same SNPs. Thus each study dataset will likely require retraining of the reference dataset. In contrast, *SNVstory* contains a pre-trained classifier that can be used as-is on any new dataset and it offers more fine-scale ancestry resolution. However, multiple models are estimated for each sample, resulting in a significantly longer run time compared to ADMIXTURE and *plinkQC*. Future improvements of the *plinkQC* ancestry module will include developing models for local ancestry estimation and training classifiers for ancestry estimation beyond continental ancestries. However, due to the small reference sample size for fine-grained populations in the current dataset, PCA projection of the external dataset will likely suffer from shrinakge to the mean, particularly for higher order PCs, so additional correction method will likely be necessary^29,30^. Lastly, to allow for custom analyses of population stratification using different reference datasets or expanding the scope to non-human genetics, we provide detailed instructions in package vignettes for how to set-up and train new models. Identifying the maximally unrelated sample set with *plinkQC* runs within minutes. While slightly longer than *King*’s runtime, *plinkQC* demonstrates larger sample retention compared to *King*’s identification process based on two estimates of relatedness.

Overall, *plinkQC* bundles together important genotype quality control functions as well as identifying related individuals and genomic ancestry identification while retaining efficiency and precision.

## Methods

### Code and Data availability

The package documentation and code are available at: https://meyer-lab-cshl.github.io/plinkQC/ and https://github.com/meyer-lab-cshl/plinkQC. Code for data analyses and manuscript figures are at: https://github.com/meyer-lab-cshl/plinkQC_manuscript. All analyses were conducted with R ≥ 4.0, plink version 1.90b6.21, and plink2 version 2.0. 0-a.6.9LM.

### Datasets

#### Combined 1000 Genomes and Human Genome Diversity Project

We downloaded the vcf files for the autosomal chromosomes of the harmonized 1000 Genomes and HGDP datasets^9^, containing 189,381,961 variants from 4,151 individuals. These variants composed of single nucleotide polymorphism (SNP), indels, and structural variants. For the purposes of ancestry identification and relatedness estimation, we merged the individual vcf files, converted the combined file to *plink* format and filtered for the 98,621,882 SNP variants only.

### Relatedness estimation

#### Data processing

From the merged dataset, we subset to samples from the 1000 Genomes project only. From these 3,202 samples, we removed ambiguous SNPs (A/T and C/G mutations) and conducted *plinkQC markerQC* to remove SNPs with high missing rate (above 1% missing) or HWE-outliers (deviation significance threshold of 1e-5).

#### Running *King* and *plinkQC*

We ran *King* on the processed data set using *king –unrelated –degree 2*. The *King* kinship score matrix was calculated using *king –kinship*. For *plinkQC* based on IBD metric, we ran *plinkQC check_relatedness()* using an IBD threshold of 0.25. For *plinkQC* based on *King* kinship, we ran *relatednessFilter ()* with a threshold of 0.0884 (second degree relative filter as defined in^5^) and supplying the *King* kinship matrix as input.

### Ancestry estimation

To prune for related individuals, we first separated the dataset into the different reference ancestral populations. We then applied *plinkQC* per-marker checks to remove markers with high missing values (above 1%), rare variants (low minor allele frequency below 0.05), and a significance threshold of 1e-05 for deviations from Hardy-Weinberg equilibrium. We removed related individuals within each ancestral label group using *plinkQC*’s check_relatedness() function repeated the marker QC as above. We pruned for genetic markers in linkage disequilibrium (LD; *r*^2^ < 0.2), with *plinkQC*’s pruning_ld() function. We then used *plinkQC*’s per-sample checks to filter out samples with outlying heterozygosity (3 standard deviations above or below the mean) and high marker missingness (above 3%). Since there were only eleven Oceania samples left after sample quality control, we removed them.

From the 3,436 samples with 229,020 SNPs that passed QC, we separated the data into 2,540 samples for the training dataset and 886 samples for the testing component. We constructed the split such that the training data included an equal number of samples from different ancestral population, when possible, based on sample size (Figure 2B, bar plot). We used *plink2 –pca* function to calculate the principal component analyses (PCA) eigenvectors and loading matrices for the training dataset. To have a consistent scale for projecting study data, we projected the training data on the loading matrices with the *plink –score* function. These projected values were used as input data to train a random forest in *R* with the *randomForest v4*.*7*.*1*.*2*^31^ and *caret v7*.*0*.*1*^32^ library.

For training, we used a grid search to optimize the number of PCs, trees, and variables sampled at each node of the random forest classifier (Supplementary Fig 1). The out-of-bag error rate for the best performing model with 11 PCs and 25 trees is 0.47%. *PlinkQC*’s ancestry identification program was run with the function: *superpop_classification()*.

#### Simulating missing data

We simulated random subsampling of a portion of SNPs from the held out testing data to represent varying levels of missing data. To do so, we randomly selected a list of SNPs and extracted them into a new dataset. We used the extracted dataset as input for the *superpop_classification()* function in *plinkQC* to determine the predicted ancestry and calculated the accuracy with the reference ancestry. We did 100 trials for each SNP percentage tested.

### Runtime Comparison

Runtime comparisons were run on a 64 bit AMD Ryzen 9 7900X 12-Core Processor Linux Ubuntu Workstation computer. Timing runs were done three times with no other program running on the computer for consistency.

#### Gambian Data Processing

FASTQ files were acquired for the Fula population from the Gambian Genome Variation Project. We used bwa-mem2^33^ to align the reads to the hg39 reference genome, followed by sorting and merging the alignments for each sample with samtools^34^. Variant calling was done with GATK HaplotypeCaller and then GenotypeGVCFs to get vcf files. Then, we filtered low-quality markers with Quality of Depth (QD) < 2.0; Fisher Strand Bias (FS) > 60.0; RMS Mapping Quality (MQ) < 40.0; Mapping Quality Rank Sum Test (MQRankSum) < −12.5; and Rank Sum Test for Relative Positioning of REF versus ALT alleles (ReadPosRankSum) < −8.0.^35^. For *plinkQC* and ADMIXTURE, this vcf file was converted into plink2 format with parameters *plink2 –vcf –not-chr X,Y,MT*. ***SNVstory*** *SNVstory* is released as a docker container that can be used with docker desktop^26^. We followed the instructions on the github (https://github.com/nch-cloud/snvstory) to install and ran *SNVstory* with parameters *–genome-ver 38 –mode WGS –sample-pos all*.

#### ADMIXTURE

Admixture version 1.3.0 was installed using conda. ADMIXTURE requires corresponding SNPs in training and study population data. Thus, we filtered the reference training dataset of combined 1000 Genomes and HGDP samples (QC for this described above) for the SNPs that are shared with the Gambian dataset before pruning variants in LD *r*^2^ < 0.2. We then trained ADMIXTURE in unsupervised mode (*–unsupervised* flag) for six random seeds and clusters were labeled manually. Visualizing the results, we observed that one trained model yielded inconsistent results, separating East Asian ancestry into two clusters and not differentiating between Middle Eastern and European ancestry, so we removed it (Supplementary Fig 3). We averaged the predicted ancestry fractions across the remaining models and then selected ids > 60% majority ancestry of the their annotated ancestry and ran ADMIXTURE on this set in supervised mode. For final ancestry prediction, we projected the Gambian genotypes using the allele frequencies for each population learned from the reference population.

## Acknowledgments

This publication uses data from the Gambian Genome Variation Project, a collaboration of the MRC Unit in the Gambia (http://www.mrc.gm), the Wellcome Sanger Institute (https://www.sanger.ac.uk), the MRC Centre for Genomics and Global Health (https://www.cggh.org/collaborations/mrc-unit-the-gambia) at the University of Oxford, and the MalariaGEN Resource Centre (https://www.malariagen.net). We also thank Audrey E. Bollas for help with troubleshooting SNVstory and Rishvanth K. Prabakar for help with processing the Gambian reference genome files.

## Supplementary Figures

**Supplementary Figure 1.**
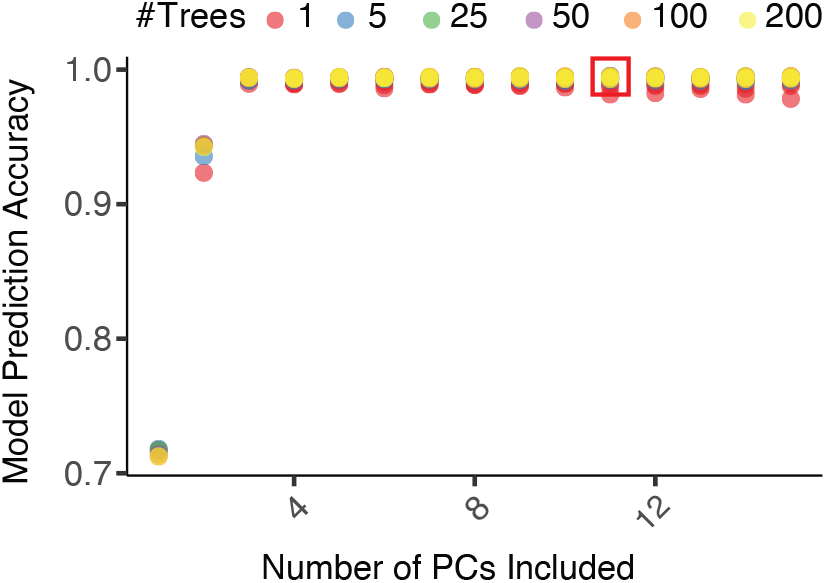
Grid Search for Optimizing Parameters. Random forest parameter selection by grid search; final parameters of classifier indicated by box.

**Supplementary Table 1.** Kinship and IBD scores for pairs of individuals labeled as related in 1000 Genomes and not removed by *plinkQC* or *King*. Individuals with related individuals (labeled by 1000 Genomes) left in the dataset after pruning to only unrelated individuals with *plinkQC* or *King*. IDs italicized and underlined remain in the dataset with a labeled relative after pruning by *plinkQC* (IBD threshold of < 0.25), *plinkQC* (kinship threshold of < 0.0884), and *King*. IDs italicized but not underlined remain in the dataset with a labeled relative after pruning by *plinkQC* (IBD threshold of < 0.25) and *plinkQC* (kinship threshold of < 0.0884). The sample HG01983 has a kinship scores with another individual within the dataset above the threshold. Thus, as described in the main text, this is another example of King removing additional samples.

| IID1 | IID2 | IBD | Kinship | Pedigree Label |
| --- | --- | --- | --- | --- |
| <u>NA12045</u> | <u>NA07031</u> | 0.0338 | -0.0034 | 2nd order |
| <u>NA19101</u> | <u>NA19092</u> | 0.1588 | 0.0624 | 2nd order |
| N19184 | <u>NA19203</u> | 0.0557 | 0.0044 | NA19203 is labeled to have N19184 as a sibling;<br>N19184 is not labeled to have NA19203 as a sibling |
| HG01936 | HG01983 | 0.2378 | 0.0775 | 2nd order |

**Supplementary Figure 2.**
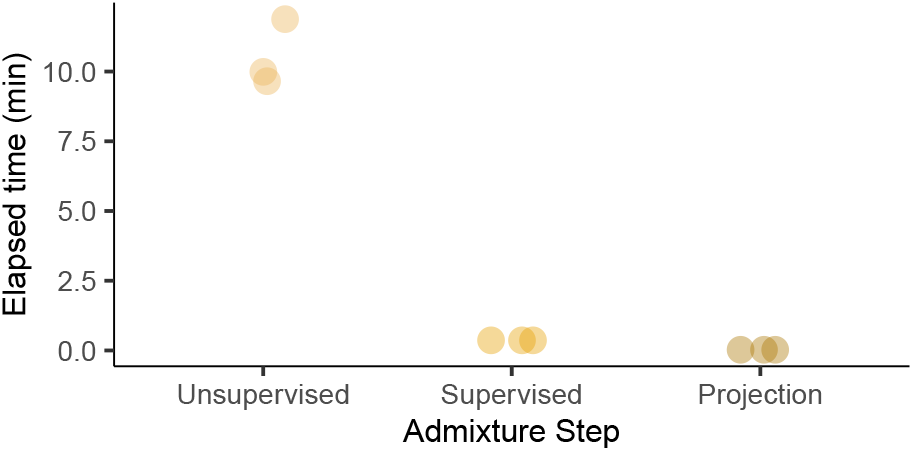
Empirical runtime analyses for ADMIXTURE. Empirical runtime analyses for the three steps (unsupervised and supervised training and classification) needed by ADMIXTURE to estimate ancestries of 100 Fulani individuals from the Gambian Genome Variation. X-axis jitter added for readability.

**Supplementary Figure 3.**
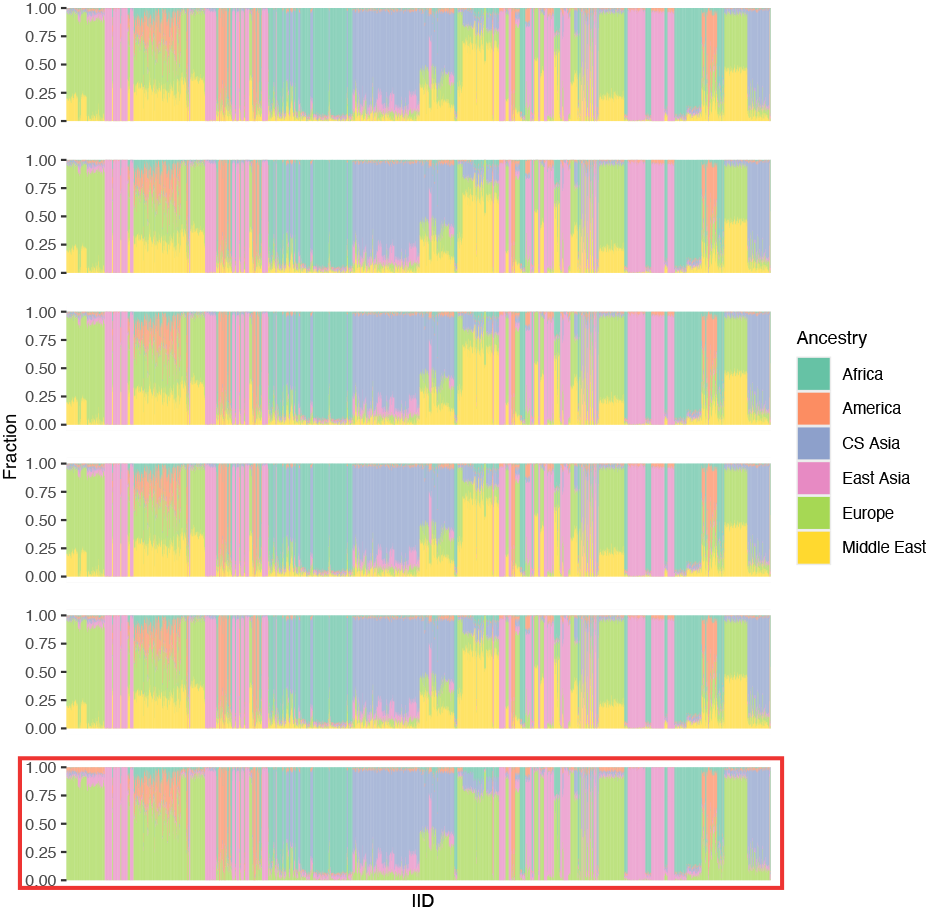
ADMIXTURE classifier after unsupervised training with six random seeds. ADMIXTURE was run in unsupervised mode with six populations on the reference dataset for six different random seeds. The model indicated with a red rectangle was removed from downstream analyses for inconsistent results compared to the remaining five models.

